# Systematically improved *in vitro* culture conditions reveal new insights into the reproductive biology of the human parasite *Schistosoma mansoni*

**DOI:** 10.1101/411397

**Authors:** Jipeng Wang, Rui Chen, James J. Collins

## Abstract

Schistosomes infect over 200 million people. The prodigious egg output of these parasites is the sole driver of pathology due to infection, yet our understanding of their sexual reproduction is limited because egg production is not sustained for more than a few days *in vitro*. Here, we describe culture conditions that support schistosome sexual development and sustained egg production *in vitro.* Female schistosomes rely on continuous pairing with male worms to fuel the maturation of their reproductive organs. Exploiting these new culture conditions, we explore the process of male-stimulated female maturation and demonstrate that physical contact with a male worm, and not insemination, is sufficient to induce female development and the production of viable parthenogenetic haploid embryos. We further report the characterization of a novel nuclear receptor, that we call vitellogenic factor 1, that is essential for female sexual development following pairing with a male worm. Taken together, these results provide a platform to study the fascinating sexual biology of these parasites on a molecular level, illuminating new strategies to control schistosome egg production.

## INTRODUCTION

Schistosomes are blood-dwelling parasitic flatworms that cause serious disease in millions of people in the developing world [1]. The pathology caused by these parasites is entirely due to the parasite’s prodigious egg output [2]. Although the goal of the parasite is to pass these eggs from the host to ensure the continuity of the parasite’s complex life cycle, approximately half of these eggs become trapped in host tissues inducing inflammation that represents the primary driver of disease [3]. Since parasites incapable of producing eggs produced little pathology in infected hosts, understanding the biology of schistosome egg production could suggest new therapeutic strategies against these devastating parasites.

Schistosomes are unique among flatworms as they do not sexually reproduce as hermaphrodites instead they have evolved separate male and female sexes [2, 4, 5]. This transition from hermaphroditism to dioecism has led to some intriguing biological phenomena, in particular the observation that female schistosomes rely on continuous pairing with a male worm to become sexually mature and produce eggs [2, 6-8]. Indeed, females grown in the absence of male worms are developmentally stunted and their reproductive organs are undeveloped. Upon pairing with a male worm, the female’s sexual organs become mature and egg production commences. Interestingly, this process is reversible since females deprived of male contact will regress to an immature state [9]. Although a variety of molecules important for female reproduction have been characterized [10-12], the nature of the signal(s) from the male that stimulate female maturation and the female response to these signals remain poorly understood.

A major bottleneck for understanding the biology of egg production and female sexual development is that normal egg production ceases within days of removal of the parasite from the host even in the presence of male worms [13-16]. While work by numerous investigators have established robust conditions for the maintenance [17-20] and growth of adult-staged parasites [15, 21, 22], no *in vitro* conditions that sustain continuous egg production have been described. Here, we report conditions that support long-term schistosome egg production *in vitro* and allow for virgin female worms to become sexually mature after pairing with a male worm. As a proof of principle, we use these culture conditions to explore the process by which male worms stimulate female maturation. We find that direct contact with a male worm along the female worm’s entire body is essential for sexual maturation and viable egg production. Interestingly, we demonstrate that in the absence of sperm transfer that contact with a male worm is sufficient for female worms to produce viable parthenogenetic haploid embryos. Capitalizing on these culture conditions we further report the characterization of a previously uncharacterized nuclear receptor that is essential for normal female development following pairing with a male worm. These studies provide new insights into the biology of schistosome egg production and lay the groundwork for the application of a growing molecular tool kit to understanding the fascinating sexual biology of these important pathogens.

## RESULTS

### Media containing ascorbic acid, red blood cells, and cholesterol supports egg production *in vitro*

The most successful systematic effort for culturing schistosomes *in vitro* are those of Basch [15, 21, 22]. While Basch’s “medium 169” (BM169) was able to support the *in vitro* growth of larval parasites to adulthood [21], it was insufficient for maintaining sexually mature egg-laying female parasites [15, 22]. Nevertheless, given the success of BM169 to support parasite growth, we reasoned BM169 was the ideal place to begin optimizing conditions for egg production. As previously reported [15], adult schistosomes recovered from mice and cultured in BM169 progressively lost the ability to lay eggs with the morphological characteristics of those laid *in vivo* or immediately *ex vivo* (Fig 1A).

**Fig 1.**
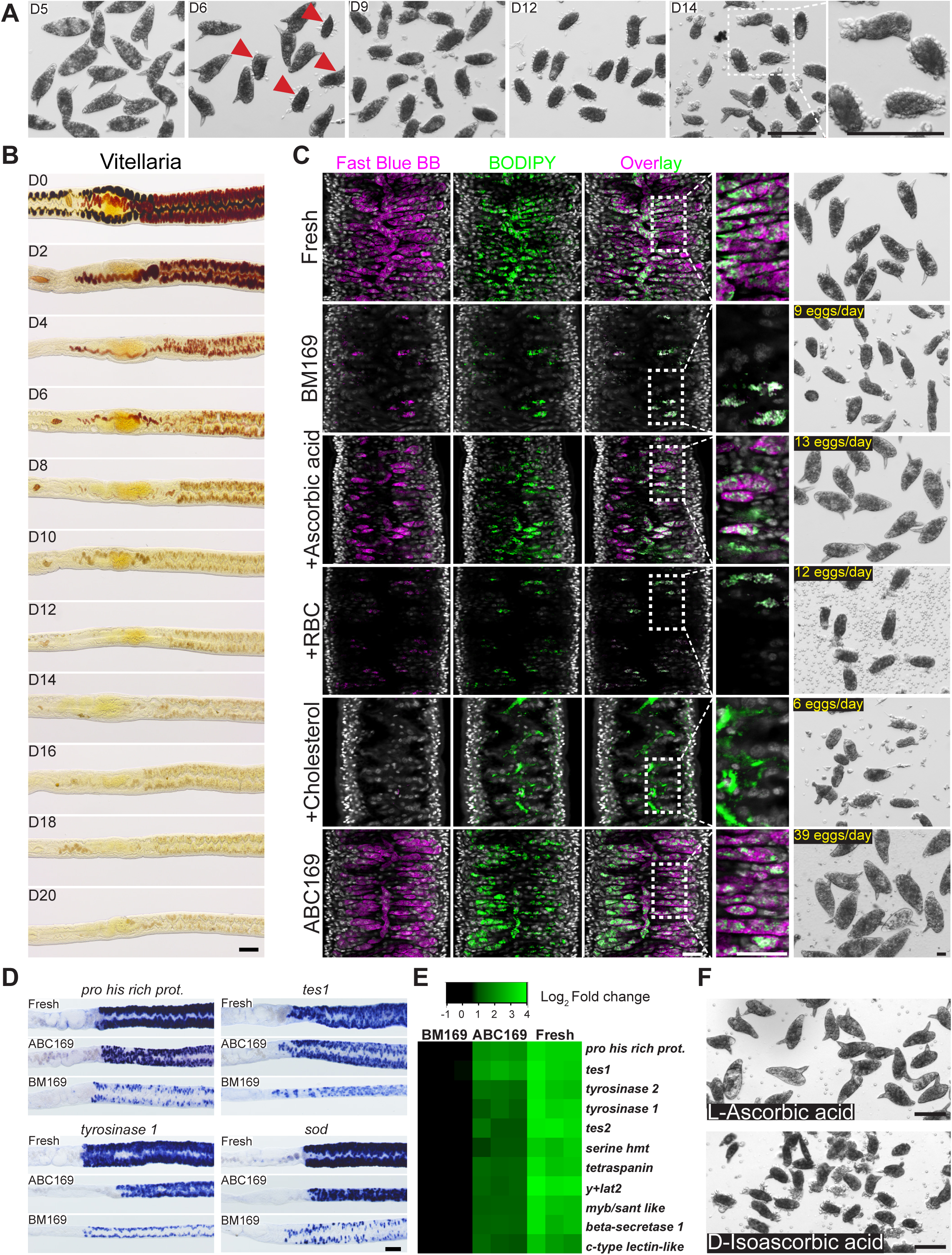
ABC169 supports the maintenance of schistosome vitellaria. (A) Morphological changes of eggs laid by worm pairs maintained in BM169. After D6 parasites began laying abnormally formed eggs (arrow). These eggs were small, usually did not contain a lateral spine, and did not possess a smooth surface. (B) Fast Blue BB staining (brownish-red labeling) showing loss of mature vitellocytes during culture in BM169. Representative images from three experiments with n > 10 parasites. (C) Confocal slice showing Fast Blue BB and BODIPY labeling in the vitellaria of freshly recovered parasites and in schistosomes at D20 of cultivation in BM169 supplemented with RBCs (Red Blood Cells), LDL, and/or ascorbic acid. Representative images from three experiments with n > 10 parasites. (D) Whole-mount *in situ* hybridization showing expression of vitellaria-enriched genes in freshly perfused female worms and parasites cultured in ABC169 or BM169 for 20 days. Representative images from three experiments with n ≥ 9 parasites. (E) Heat-map showing relative expression of vitellaria-enriched transcripts in freshly perfused female worms (“Fresh”) and parasites cultured in ABC169 or BM169 for 20 days. Each column represents an independent biological replicate; samples are normalized to the expression of an arbitrarily chosen biological replicate from the BM169 group. Changes in expression between BM169 and ABC169 were statistically significant (p<0.05, T-test). (F) Morphology of eggs laid by paired adult females in medium ABC169 supplemented with L-ascorbic acid or D-isoascorbic acid on D20. Representative of 3 experiments. Scale bars: A, B, D, F 100µm; C 25µm.

A schistosome egg is constructed from cells derived from two organs: the ovary, that contributes an oocyte, and the vitellaria, that provide 20-30 vitellocytes [2, 23]. Although female worms paired with male worms retained the ability to produce oocytes during *in vitro* culture (Fig S1A,B), we noted a rapid loss in the ability of cultured parasites to generate vitellocytes, consistent to previous studies [14, 16]. Vitellocytes contain two types of large cytoplasmic inclusions: lipid droplets and vitelline droplets that coalesce to form the eggshell [24, 25]. Using Fast Blue BB to label vitelline droplets we found that vitellaria progressively ceased production of large numbers of vitellocytes in BM169 (Fig 1B,C). Similarly, BODIPY 493/503 labeling found that the vitellaria of females in BM169 possessed few lipid droplets at D20 (Fig 1C). Examination of a panel of genes expressed in mature vitellocytes [26] by whole mount *in situ* hybridization (Fig 1D) and quantitative PCR (Fig 1E) found a significant decrease in the expression of vitellaria-specific transcripts during culture, similar to previous studies [16]. Thus, the capacity for vitellogenesis is rapidly lost *in vitro*.

To improve the rate of vitellogenesis and egg production we examined supplements that could potentially satisfy either known metabolic requirements for egg production (e.g., lipids[27]) or documented auxotropies of the worm (e.g., polyamines, fatty acids, sterols[28-30]). From these analyses we observed no qualitative effects on egg production from supplements, including: albumin (e.g., Lactalbumin or Linoleic Acid-Oleic Acid-Albumin), spermidine, commercially available lipid supplements, commercially available antioxidant supplements, red blood cells (RBCs), low density lipoprotein, L-carnitine, N-acetyl-cysteine, or sera from various species (chicken, bovine, or horse). During this process, however, we came across a report detailing the formation of abnormal eggs in schistosome-infected guinea pigs fed an Vitamin C (L-ascorbic acid) deficient diet [31]. Strikingly, addition of ascorbic acid to BM169 led to a marked increase in vitelline development (Fig 1C, S1C Fig) and the production of eggs morphologically similar to those laid by parasites immediately *ex vivo* (Fig 1C).

Although L-ascorbic acid had profound effects on the quality of eggs generated *in vitro*, the rate of egg production and development of the vitellaria remained inferior to that of fresh *ex vivo* parasites (Fig 1C). Thus, we re-examined some of the previous assessed media supplements. Given the critical role for lipid metabolism in egg production [27], we reasoned that adding complex sources of lipids (and other nutrients) that the parasite encounters *in vivo* might act synergistically with ascorbic acid. We found that supplementation with either red blood cells or a commercial “cholesterol” concentrate containing purified Low Density Lipoprotein (LDL) increased lipid stores along the intestine but had little effect on vitelline development or the production of normal eggs (Fig 1C). However, combination of red blood cells, the cholesterol/LDL concentrate, and ascorbic acid produced a dramatic increase in the rate of vitellogenesis (Fig 1C), the rate of egg production (Fig 1C), and a marked increase in the expression of vitellaria-specific transcripts (Fig 1D,E). From here on we refer to this formulation as ABC169 (<u>A</u>scorbic Acid, <u>B</u>lood Cells, <u>C</u>holesterol and Basch Media <u>169</u>).

In vertebrate cells, L-ascorbic acid acts not only as an antioxidant, but an essential co-factor for a variety of enzymes [32]. Mammals deficient for L-ascorbic acid cannot perform key enzymatic reactions (e.g., collagen hydroxylation) leading to symptoms commonly known as scurvy [32]. Interestingly, the effects of vitamin C to prevent scurvy are stereoselective as D-isoascorbic acid cannot replace L-ascorbic acid at equimolar concentrations [33]. Similarly, we observed that D-isoascorbic acid could not replace L-ascorbic acid in egg production (Fig 1F), suggesting that either L-ascorbic acid is selectively transported into cells or that it acts in a stereoselective fashion to facilitate one or more enzymatic reactions within the cell.

### Parasites cultured in ABC169 produce eggs capable of developing to miracidia

After an initial peak, egg production in BM169 dropped precipitously and by D7 of culture parasites laid ∼13 egg-like masses per day (Fig 2A,B). We noted a similar peak in egg production using ABC169, however, after D7 these parasites sustained production of ∼44 morphologically normal eggs per day (Fig 2A). Indeed, eggs laid in ABC169 possessed a lateral spine typical of *S. mansoni* eggs, had smooth shells, and contained a “germinal disc” corresponding to the early embryo (Fig 2B). Eggs freshly laid in ABC169 media were larger on average than those laid in BM169 (Fig 2C) and contained similar numbers of nuclei as eggs laid by parasites freshly recovered from mice (Fig 2D). Due to a high concentration of phenolic proteins that originate in the vitelline droplets that form the eggshell, schistosome eggshells are highly autofluorescent [23]. In contrast to the egg-like masses from BM169, where parasites produce few vitelline droplets (Fig 1C), eggs from parasites in ABC169 possessed autofluorescence comparable to those from parasites freshly *ex vivo* (Fig 2E). Furthermore, nearly half of the eggs laid by parasites at D18-20 of culture in ABC169 contained clusters of proliferative embryonic cells visualized by labeling with thymidine analog EdU (Fig 2F, G). *S. mansoni* eggs are passed from the host to release larvae called miracidia [2]. Approximately 10-20% of eggs laid on the first day cultured either in BM169 or ABC169 produced miracidia (S2 Fig). However, this rate dropped during time in culture (S2 Fig). While eggs laid in BM169 after D7 were incapable of liberating miracidia, about 1-2% of eggs laid in ABC169 produced viable and morphologically normal miracidia (Fig 2H, S1-2 Movie). Furthermore, we found that miracidia recovered from eggs laid on D15 to D20 were capable of penetrating *Biomphalaria glabrata* snails (n=214/217 total miracidia). However, after examining more than 25 infected *B. glabrata* snails we only found two snails that shed viable cercariae (S3 Movie). Given the disparity between the rate of entering embryonic development (Fig 2G) and the capacity for these embryos to mature to miracidia (S2 Fig), together with the limited ability of these miracidia to generate cercariae after infecting snails, it is possible that cues from the host are necessary for efficient development of *S. mansoni* embryos to infective miracidia. Further studies exploring optimal growth conditions for schistosome egg development could address these issues.

**Fig 2.**
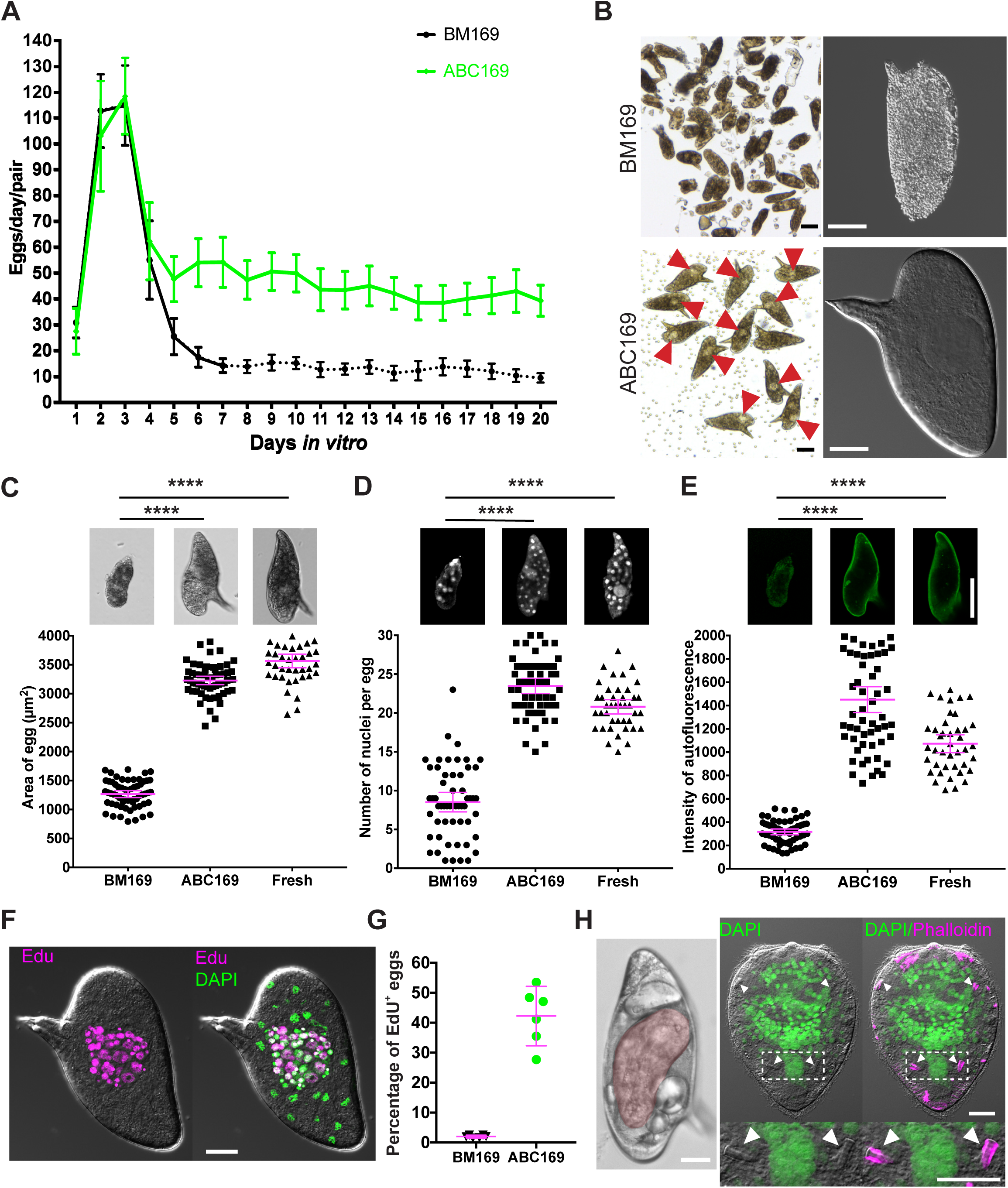
Eggs laid in ABC169 develop and hatch. (A) Rate of egg production per worm pair in BM169 or medium ABC169 during *in vitro* culture. Dotted line indicates the point at which parasites beginning laying morphologically abnormal eggs. n ≥ 19 worm pairs examined in 4 experiments. Error bars represent 95% confidence intervals. (B) Morphology of eggs (Left, bright field; Right, DIC) laid by paired adult females in BM169 or medium ABC169 on D20. Red arrows show early embryos in eggs laid in ABC169. (C-E) Quantification of the (C) size, (D) number of DAPI-labeled nuclei, and (E) autofluorescence intensity from eggs laid by freshly perfused female worms (Fresh, n=41), parasites cultured in BM169 (n=61), or ABC169 (n=59) on D20. **** p<0.0001, T-Test. Error bars represent 95% confidence intervals. (F) EdU-labeled embryonic cells of an egg laid by a paired adult female in ABC169 on D20. (G) Percentage of eggs with clusters of cycling EdU^+^ embryonic cells from eggs laid between D16 and D22 by paired adult females in BM169 or ABC169. >2000 eggs from 6 experiments. Error bars represent 95% confidence interval. (H) Miracidia from eggs laid in ABC169. Left, miracidium (pseudocolored red) inside an egg laid on D20 of culture in ABC169. Right, hatched miracidium from D20 egg labeled with DAPI and phalloidin. These miracidia appear grossly normal in morphology possessing 2 pairs of flame cells (arrows). Scale bars: B, F, H 20 µm; C-E 50 µm.

### Immature virgin females sexually mature and produce viable eggs upon pairing with male worms in ABC169 medium

Studying the process by which male schistosomes stimulate female development has traditionally been challenging since females experience incomplete development following pairing with a male *in vitro* [34, 35]. To examine sexual maturation in ABC169, we recovered immature virgin females from mice and paired these females with sexually mature virgin male worms. The vitellaria of paired immature females grown in BM169 were poorly developed (Fig 3A) and these parasites laid small numbers of morphologically abnormal eggs (Fig 3B). However, the vitellaria of newly paired immature virgin females cultured in ABC169 developed normal vitellaria (Fig 3A) and these parasites were capable of laying morphologically normal eggs beginning between D6-D7 of culture (Fig 3B-C). These eggs could initiate embryogenesis (Fig 3D, 42.7% n=1010 from D16 to D22) and about 2% could develop to miracidia (eggs from D11-14, n=2807 eggs). Thus, ABC169 supports female development following pairing with a male worm.

**Fig 3.**
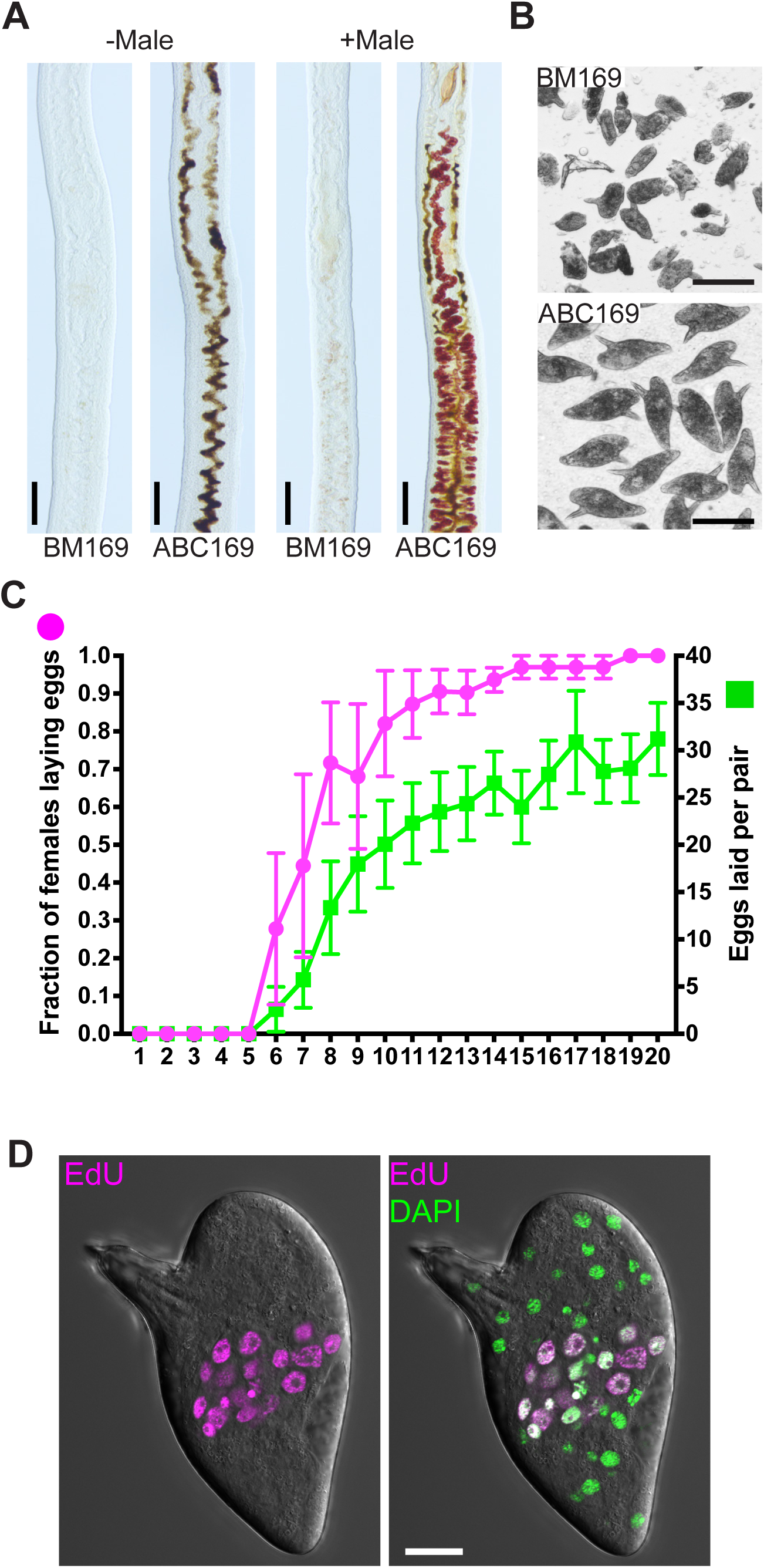
ABC169 supports the maturation of immature female schistosomes following pairing with a male worm. (A) Vitellaria development visualized by Fast Blue BB labeling in immature female schistosomes in the presence or absence of male worms in BM169 or ABC169 at D20 of culture. Black pigment in ABC169 parasites represents digested RBCs. (B) Eggs laid by immature female in BM169 or ABC169 on day 20 after pairing with a male. (C) Proportion of females laying eggs (left axis, magenta) and number of eggs laid per worm pair (right axis, green) from D1 to D20 following pairing of immature females with male worms in ABC169. n=31 worm pairs examined in 4 separate experiments. Error bars represent 95% confidence intervals. (D) Eggs laid by previously immature females commence embryonic development. Left, representative developing EdU-labeled embryo from egg laid in ABC169 on D20. Scale bars: A,B 100 µm; D 20 µm.

### Physical contact with a male worm is sufficient for female worms to produce viable parthenogenetic haploid embryos

Several theories have been put forward to explain the mechanism by which male worms stimulate female development [2, 36-38]. However, the experiments supporting (or refuting) these hypotheses were conducted using sub-optimal culture conditions [37] and in many cases have not been subject to extensive reproduction in the modern literature. Thus, we were compelled to revisit key observations using ABC169. The prevailing thought is that female development requires direct contact with a male worm [7, 37, 38]. The most intriguing study supporting this hypothesis are those of Popiel and Basch [35]. These authors observed that small segments of male worms could stimulate vitelline development. Interestingly, vitelline development was confined to regions in direct contact with the male segment [35]. Consistent with these observations, we found a large fraction of small posterior fragments could pair with immature females (Fig 4A,B, S4 Movie). These posterior segments often paired with posterior regions of female worms, and consistent with observations of Popiel and Basch, vitelline maturation occurred only in regions in direct contact the male segment (Fig 4B). Thus, pairing with a male induces a signal that induces localized female vitelline maturation.

**Fig 4.**
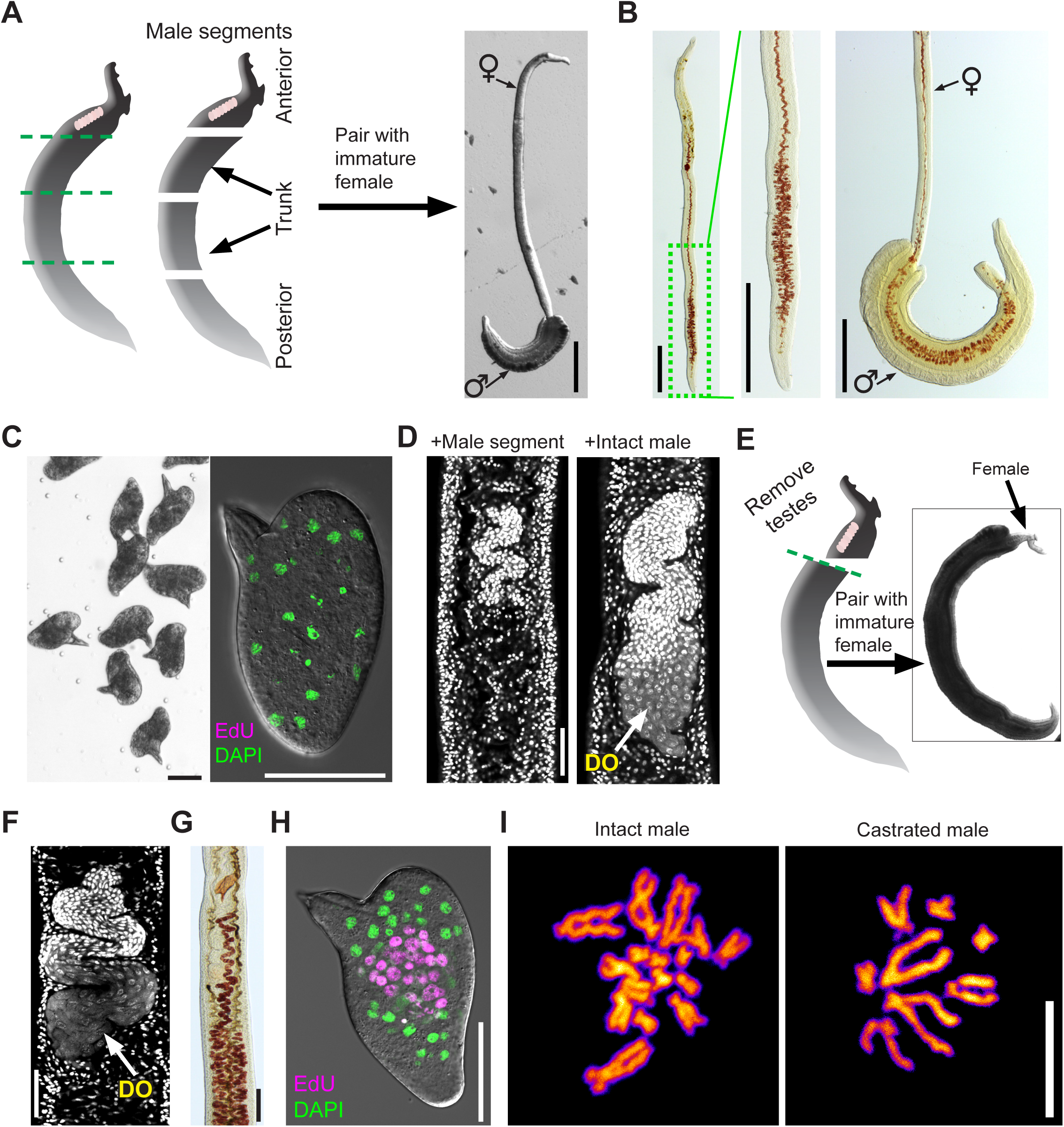
Using ABC169 to study schistosome male-induced female sexual maturation. (A) Pairing of immature females with amputated male segments. Left, cartoon showing approximate positions of male amputations. Most often we observed posterior segments pairing with posterior region of immature females, shown to right. (B) Fast Blue BB labeling showing female vitellaria development after pairing with posterior male segments. Left panels, female after separation from male segment showing vitellaria development in posterior region. Right panel, female before separation from male segment showing vitellaria development is restricted to paired region. Representative of > 40 female parasites examined in 4 separate experiments. (C) Eggs laid by immature female worms paired with male segments are morphologically normal (left) but do not contain developing embryos as measured by EdU labeling (right). n > 500 eggs examined in 4 separate experiments. (D) DAPI staining to examine ovary development in immature females paired with intact male worms or an amputated posterior male segment. Females paired with intact males possess differentiated oocytes (DO) (n=23), whereas females paired with male segments produce no differentiated oocytes (n=41). (E) Pairing of immature females with decapitated and castrated male segments. Left, cartoon showing approximate positions of amputation, removing both the head and testes. These decapitated and castrated segments paired with female along most of the female body, shown to right. (F) DAPI staining to examine ovary development in immature females paired with decapitated and castrated male segments. These ovaries contain differentiated oocytes (DO). Representative images from n = 22 parasites examined in 3 separate experiments. (G) Fast Blue BB showing vitellaria development in females after pairing with decapitated and castrated male segments. Representative images n=22 parasites from 3 separate experiments. (H) EdU labeling showing embryonic development in eggs from females paired with decapitated and castrated male segments. 273/602 eggs laid between D16-D22 contained clusters of EdU^+^ cells. (I) DAPI labeling of metaphase spreads from eggs laid by (left) fresh *ex vivo* female parasites paired with intact males and (right) immature female parasites paired with decapitated and castrated male segments. Embryonic cells from fresh *ex vivo* parasites were diploid (2n=16), whereas those from unfertilized females are haploid (n=8). Scale bars: A, B 500 µm; C, D, F, H 50 µm, G 100 µm, I 10 µm.

While culturing male posterior segments with female worms we observed that these female worms laid morphologically normal eggs (Fig 4C). However, examination of these eggs found that they contained no embryos capable of incorporating EdU (Fig 4C). To explore this observation in more detail we examined the ovaries of the female worms paired with male segments and found no evidence of mature oocyte production (Fig 4D). Since the schistosome ovary is located anterior to the vitellaria, and this region was not in contact with male posterior segments, we reasoned that ovaries, like the vitellaria, might also require local contact with a male worm to mature and begin oocyte production. To test this model, we amputated males behind the testes (Fig 4E) and paired the decapitated and castrated fragments with immature female worms. Since these large posterior fragments typically ensheathed the entire female worm (Fig 4E), we reasoned that this pairing might be sufficient to stimulate oogenesis. Consistent with this model, we found that ovaries of females paired with decapitated males produced oocytes (Fig 4F). Furthermore, these parasites had fully developed vitellaria along their entire length (Fig 4G) and laid morphologically normal eggs (Fig 4H).

Despite the fact that females paired with castrated male segments had no chance of being inseminated, we observed that eggs laid by these parasites possessed the ability to initiate embryogenesis (Fig 4H) and could even give rise to miracidia (0.23%, n=4704 eggs). Previous studies have suggested that female schistosomes can produce parthenogenetic offspring containing a haploid set of maternal chromosomes when mated with males of distantly related schistosome species [39, 40]. From these experiments it is not clear if simply coming into contact with a male schistosome of another species is sufficient to induce the production of parthenogenetic offspring or if parthenogenesis occurs only following the transfer of sperm [39, 40]. Therefore, we examined the karyotypes of eggs laid by females paired with castrated males. Unlike the diploid karyotypes of embryos from eggs laid by freshly *ex vivo* parasites (2n=16), mitotic cells from eggs laid by females paired with castrated males were haploid containing only 8 chromosomes (Fig 4I, S3 Fig). These results suggest that only contact with a male worm along the entire length of the female body, and not insemination, is sufficient for the production of viable haploid embryos. These data highlight the value of ABC169 media to enhance our understanding of schistosome reproductive biology.

### A novel nuclear receptor is required for the maturation of female vitellaria following pairing with a male worm

The ability of virgin females to mature upon pairing with a male in ABC169 (Fig 3A) provides an opportunity to discover novel regulators of male induced female maturation. In virgin female worms, specialized stem cells located in primordial ovaries and vitellaria differentiate to oocytes or vitellocytes, respectively, after paring with a male worm. Thus, the stem cells in the primordial reproductive organs must be capable of responding to external cues instructing these cells to produce differentiated progeny (i.e., oocytes or vitellocytes). Therefore, characterizing genes expressed in these stem cells could provide clues about molecules essential for maturation following pairing.

To discover genes expressed in the reproductive stem cells, we queried our previously published dataset of genes expressed in the mature female vitellaria [26] and performed *in situ* hybridization on mature and immature female worms. From this analysis, we identified *Smp_248100* (previously known as *Smp_212440*), an uncharacterized protein that shares similarity to members of the nuclear receptor family of ligand-activated transcription factors. Canonical nuclear receptors contain two key domains: a highly-conserved N-terminal DNA-binding domain (DBD) and a C-terminal ligand-binding domain (LBD)[41]. Primary sequence analysis confirmed that Smp_248100 contained a DBD with high amino acid identity to DBDs from other vertebrate and invertebrate nuclear receptors (Fig 5A and S4A Fig). However, examination of the Smp_248100 C-terminus failed to identify a canonical LBD. Interestingly, the C-terminus of Smp_248100 shared stretches of high amino acid identity with orthologous proteins from other parasitic flatworms (S4B Fig), leaving open the possibility that this protein contains a divergent LBD.

**Fig. 5.**
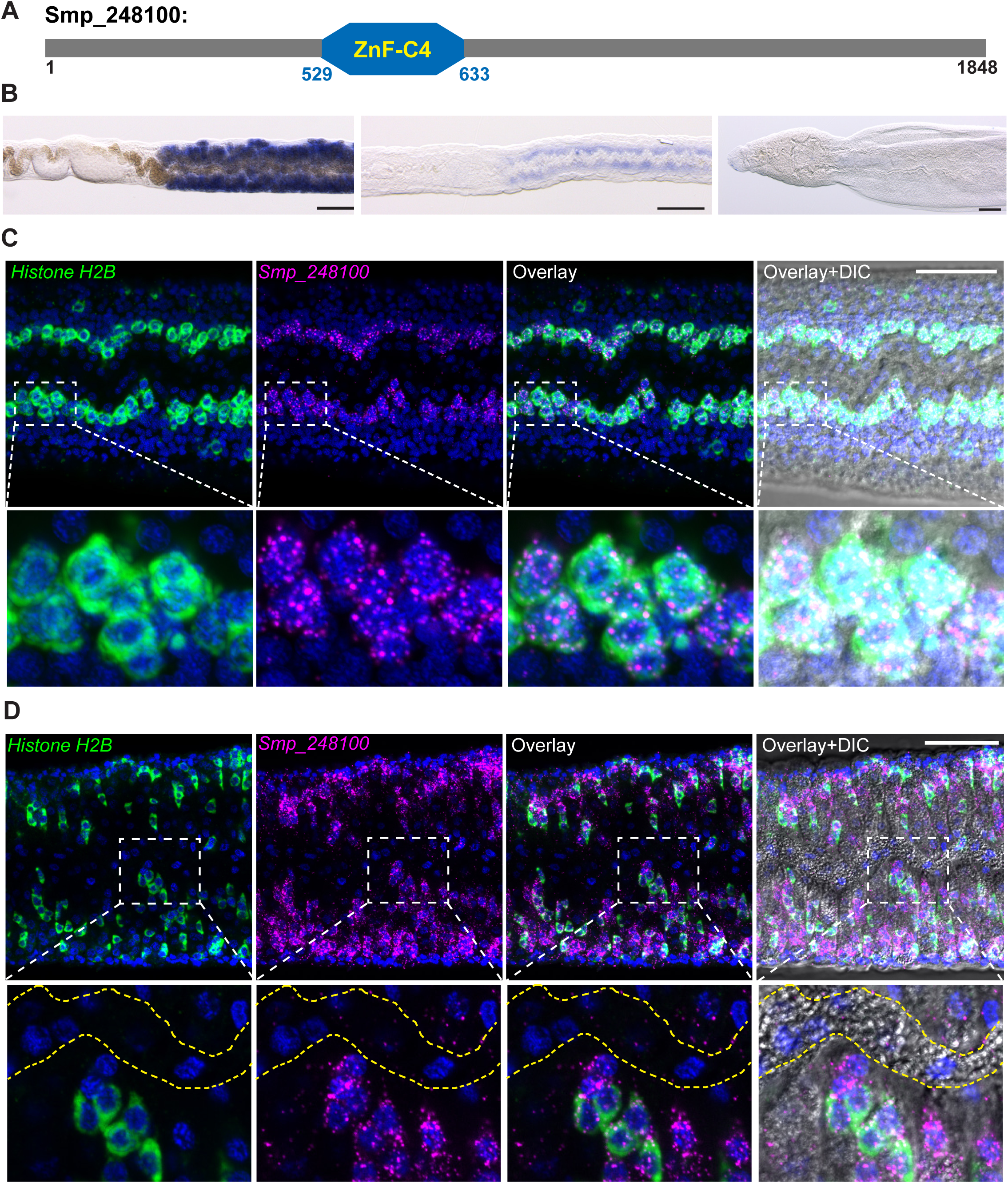
*Smp_248100* is expressed in the primordial vitellaria. (A) A schematic diagram of protein product encoded by *Smp_248100*. This protein contains a ZnF-C4 domain similar to other nuclear receptor proteins. (B) Whole-mount *in situ* hybridization showing expression of *Smp_248100* in a mature female (left), immature female (middle) or male worm (right). *Smp_248100* highly expressed in female worms exclusively in the region of vitellaria. Representative images from three experiments with n >35 parasites. Scale bar 100 µm. (C-D). Two-color FISH for *Histone H2B* (green) and *Smp_248100* (magenta) mRNAs on (C) immature females or (D) sexually mature females. In immature virgin females, *Smp_248100* was only expressed in *H2B*^+^ cells. In adult female, *Smp_248100* was expressed in *H2B*^+^ cells as well as some *H2B*^-^ cells. We did not detect *Smp_248100* expression in mature vitellocytes in the vitelline duct (bounded by dashed yellow lines). Representative images from n >30 parasites examined in 3 separate experiments. Scale bars: B, 100 µm; C, 50 µm.

By colorimetric *in situ* hybridization, we detected *Smp_248100* expression exclusively in the vitellaria of mature and immature female (Fig 5B). We failed to detect high-levels of specific expression in male parasites (Fig 5B). In immature females, the proliferative stem cells of the primordial vitellaria, also known as S1 cells [24], lie along the intestine in the posterior of the worm [16]. Thus, based on the expression of *Smp_248100* in immature females we reasoned that this gene is expressed in the S1 stem cells of the primordial vitellaria. To examine this model, we used a proliferative cell marker, *Histone H2B*, to label the proliferative S1 cells [26]. Double fluorescent *in situ* hybridization (FISH) on immature female worms found that most *H2B* ^+^ cells along the gut also expressed *Smp_248100* (Fig 5C), indicating *Smp_248100* is expressed in S1 cells within the primordial vitellaria. In mature females, we similarly detected *Smp_248100* in *H2B*^*+*^ S1 cells, however, we also detected *Smp_248100* expression in *H2B*^*-*^ cells adjacent to *H2B*^*+*^ cells (Fig 5D). Since we did not detect *Smp_248100* expression in fully mature vitellocytes with in the vitelline duct (Fig 5D), we suggest that *Smp_248100*^*+*^ *H2B*^*-*^ cells are likely the immediate differentiation progeny of the S1 cells.

Based on the expression pattern, we hypothesized that *Smp_248100* may play a role in vitelline cell development. To examine this possibility, we performed RNA interference (RNAi) on immature virgin female worms in ABC169 and examined whether these parasites could become sexually mature and commence egg-laying upon pairing with male worm. We observed majority of the *Smp_248100(RNAi)*-treated female parasites failed to produce mature vitellocytes as measured by Fast Blue BB staining (Fig 6A, B, S5 Fig) and *in situ* hybridization using the vitellocyte marker *superoxide dismutase* [26] (Fig 6B). Consistent with the effects of *Smp_248100* RNAi treatment on vitelline cell development, we noted a significant decline in the rate of egg production compared to controls (Fig 6C). Since *Smp_248100* is expressed in the proliferative S1 cells, we reasoned that the *Smp_248100(RNAi)* may result in either defects stem cell maintenance or a defect in the ability of these cell to differentiate. To distinguish between these possibilities, we examined cell proliferation in the region of the vitellaria following *Smp_248100* dsRNA treatment. We observed no difference in cell proliferation in the region of the vitellaria in *Smp_248100(RNAi)* parasites (Fig 6D), suggesting that these gene is dispensable for S1 cell maintenance. Thus, *Smp_248100* appears to be a key regulator of stem cell differentiation in the schistosome vitellaria. We also examined the effects of *Smp_248100* dsRNA treatment on ovary development. Consistent with a lack of robust *Smp_248100* expression in the ovary, we detected no defect in the ability of *Smp_248100* RNAi-treated parasites to produce differentiated oocytes (Fig 6E). Given this specific role for Smp_248100 in vitelline cell development we propose this protein be called Vitellogenic Factor 1 (VF1).

**Fig. 6.**
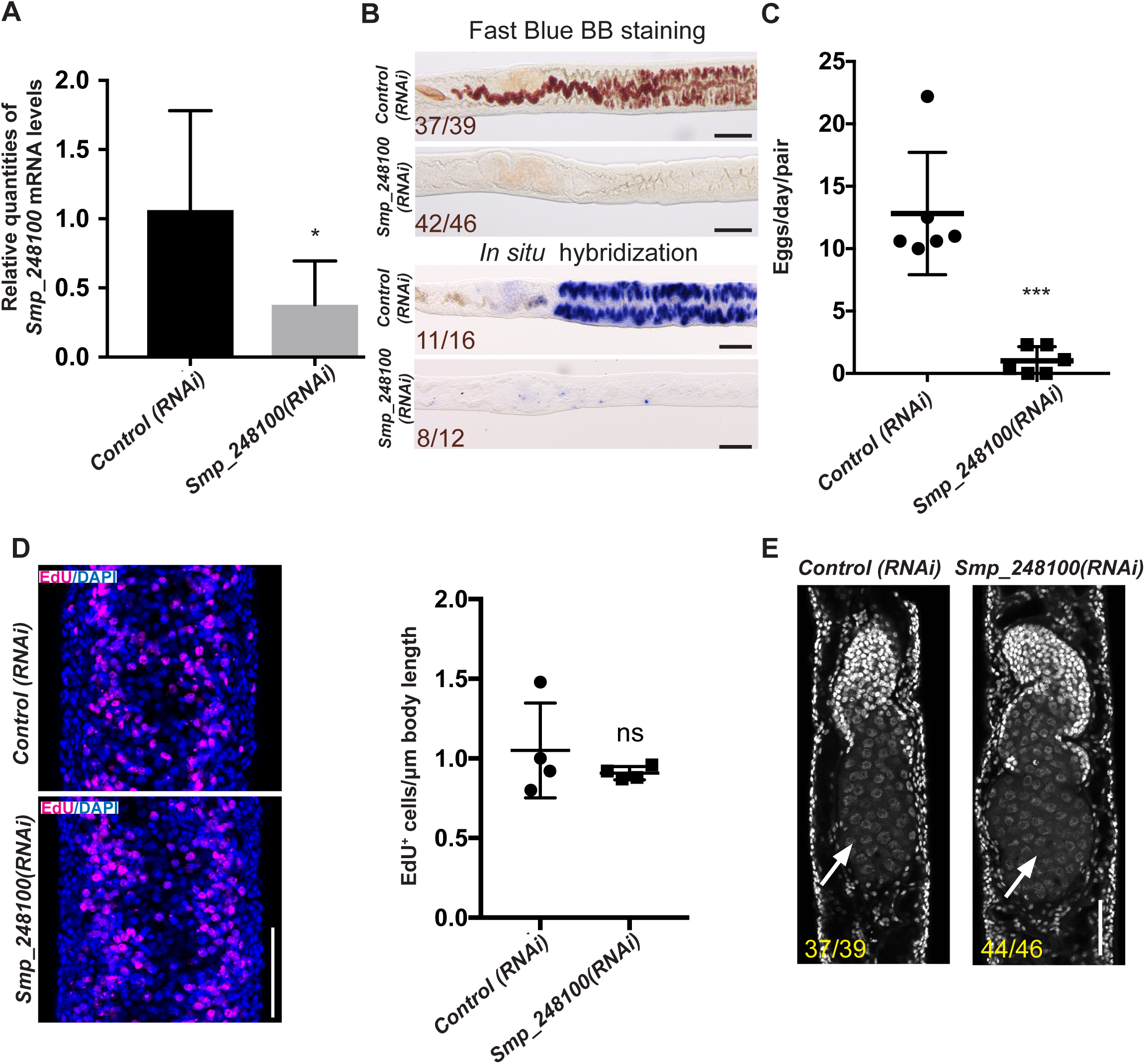
*Smp_248100* is specifically required for vitellaria maturation following pairing with a male worm. (A) qPCR showing decreased *Smp_248100* transcript levels in *Smp_248100(RNAi)* versus *control(RNAi)*. * p<0.05. Error bars indicate 95% confidence intervals calculated based on 4 separate experiments. (B) Loss of mature vitellaria in *Smp_248100(RNAi)* worms. Fast Blue BB labeling following control or *Smp_248100* RNAi treatment (top). Whole-mount *in situ* hybridization showing expression of *sod* gene (a marker for vitellocytes) in control RNAi or *Smp_248100(RNAi)* worms (bottom). *Smp_248100(RNAi)* parasites produce few vitellocytes. Representative images from 4 separate experiments. (C) Plot showing egg production per worm pair between D12-D14 of RNAi treatment. Representative images from n > 146 parasites for each treatment examined in 6 separate experiments. *** p<0.001, T-Test. Error bars represent 95% confidence intervals. (D) EdU labeling of proliferative cells following control or *Smp_248100* RNAi treatment. We observed no significant difference in the number of EdU-labeled nuclei between *control(RNAi)* and *Smp_248100(RNAi)* animals within the vitellaria. Representative images from n > 41 parasites for each treatment examined in 4 separate experiments. ns, p>0.05, T-Test. Error bars represent 95% confidence intervals. (E) DAPI labeling showing ovaries from control and *Smp_248100* RNAi-treated female parasites at D14 after pairing with a male worm. Anterior towards the top, arrow indicates differentiated oocytes. Representative images from 4 separate experiments. Scale bars: B, 100 µm; D,E 50 µm.

## DISCUSSION

Here we describe culture conditions that support sustained schistosome sexual development i*n vitro*. Of all the components of ABC media, L-ascorbic acid appeared to have the most profound effect sexual development and egg production. Since the effects of L-ascorbic acid are stereoselective (Fig 1F), it is likely its effects are related to either the transport of L-ascorbic acid into the cell or its activity on specific biological molecules. One potential target of L-ascorbic acid is the tyrosinase enzymes present in mature vitellocytes. During the production of schistosome egg shells, tyrosinases in the vitellocytes are essential for crosslinking eggshell proteins that coalesce to form the eggshell [42, 43]. *In vitro* studies on tyrosinases suggest that L-ascorbic acid can enhance the activity of tyrosinases by reducing copper molecules in the active site of the enzyme [44]. Consistent with the model that L-ascorbic acid potentiates schistosome tyrosinase activity, inhibition of tyrosinase activity in female schistosomes results in eggshell deformations [43] that appear similar to eggs from schistosomes grown *in vitro* without L-ascorbic acid (Fig 1A). Future studies aimed at determining the function of vitamin C in schistosomes could suggest new approaches to blunt egg production.

Our studies confirmed the importance direct physical contact with a male in inducing female sexual maturation. Indeed, we found that contact with a castrated male along the entire length of the female body was sufficient to induce sexual maturation and the production of parthenogenetic haploid embryos (Fig 4I). The phenomenon of parthenogenetic sexual reproduction in *Schistosoma mansoni* has been observed several times in studies performing intraspecies crosses (e.g., female *S. mansoni* x male *S. japonicum*) inside mammal hosts [39, 40, 45]. However, given the propensity of certain schistosome species to form intraspecies hybrids [46], it is challenging to distinguish between genetic hybridization and true parthenogenesis. Since our studies were conducted entirely under *in vitro* conditions using virgin females and castrated males we believe these are the strongest evidence to date that female schistosomes require only contact with a male worm to generate viable haploid progeny. As studies in the field continue to explore the genetics of schistosomes in regions endemic for multiple schistosome species [47], it will be interesting to see if the production of parthenogenetic haploid offspring is common in natural schistosome populations.

Here we demonstrate an essential role for the nuclear receptor VF1 in the differentiation of schistosome vitellocytes. Several nuclear receptors (e.g, the estrogen receptor) are activated by sex hormones and go on to elicit transcriptional responses key to sexual development [48]. Given this, it is tantalizing to speculate that that VF1 executes a pro-vitellogenic transcriptional program in response to activation by a schistosome sex hormone. Since our analyses sequence failed to identify a canonical LBD in the C-terminus of VF1, it is possible that VF1 lacks the capacity to bind traditional hormone receptor ligands (e.g., steroids). However, the possibility remains that VF1 is activated by a non-canonical nuclear receptor ligand. Indeed, examination of VF1 orthologs in other parasitic flatworms uncovered a region of extensive amino acid identity in the C-termini of these proteins (S4B Fig). Since these other flatworms also possess vitellaria, and the evolution of vitellaria is a relatively recent innovation in flatworm biology [4, 49], it is possible that during vitellaria evolution these VF1 proteins adapted to being activated by a non-canonical ligand. In addition to their ability bind ligands, nuclear receptors are notorious for their propensity to bind DNA as either homo-or heterodimers [41]. Thus, VF1 may lack ligand-binding activity but retain the ability to heterodimerize with other ligand-activated nuclear receptors. One set of potential heterodimerization partners for VF1 are the schistosome RXR nuclear receptors. Indeed, RXR proteins heterodimerize with the variety of NRs [41] and the two schistosome RXR proteins are known to bind the promoters of genes encoding egg shell proteins in the vitellaria [50, 51]. Thus, there is the possibility these nuclear receptors collaborate to coordinate pro-vitellogenic cell fates.

What role might VF1 play in male-induced female maturation? Clearly VF1 is essential for vitelline cell development following pairing with a male worm. However, we did not detect *vf1* expression in the ovary and our RNAi experiments suggest that oogenesis is not affected in *vf1(RNAi)* parasites. Thus, VF1 is not likely to be a global regulator in the female response to pairing. A more likely scenario is that VF1 is activated by pairing, resulting in transcriptional changes important to vitellogenesis in the primordial vitellaria. Based on this model, we would also speculate there are ovary-specific transcriptional regulators that are activated upon pairing within the primordial ovary. Capitalizing on ABC media, and modern molecular approaches, future studies should be aimed at uncovering the general upstream signaling programs that activate factors, such as VF1, in the primordial reproductive organs.

During his extensive studies to develop *in vitro* culture conditions for *S. mansoni*, Paul Basch lamented that the production of viable eggs *in vitro* “remains a formidable challenge” [22]. We find three additions to Basch’s base media formulation (ascorbic acid, blood cells, and LDL) can sustain the production of viable eggs for several weeks *in vitro*. Although ABC169 does not fully replicate the *in vivo* reproductive potential of the parasite, it allows us to recapitulate important aspects of schistosome sexual biology *in vitro* and thus represents a significant improvement over existing culture methods. Taken together, the ability to sustain sexual maturity *in vitro,* coupled with a growing understanding of the molecular programs associated with distinct reproductive states [52, 53], and tools such as RNAi, will expedite our understanding of the molecular factors governing parasite sexual development and egg production, perhaps suggesting new therapeutic opportunities.

## Materials and Methods

### Parasites and *in vitro* maintenance

Adult NMRI *Schistosoma mansoni* (6-7 weeks post infection) were recovered from infected mice by perfusion through hepatic portal vein with 37°C DMEM (Mediatech, Manassas, VA) plus 5% serum and heparin (200–350 U/ml). Single sex infections were obtained by infecting mice with male or female cercariae recovered from NMRI *Biomphalaria glabrata* snails infected with single miracidia. Following recovery from the host, worms were rinsed several times in DMEM + 5% serum before placing into culture media. The base medium for these studies was BM169 [21] with the addition of 1x Antibiotic-Antimycotic (Gibco/Life Technologies, Carlsbad, CA). Although Basch included red blood cells in BM169, it has been noted that erythrocytes, which schistosomes consume *in vivo*, are not sufficient to sustain egg production [16, 22, 27] and were initially omitted from the base medium. We explored sera from a variety of sources (horse, fetal bovine, chicken) and determined that newborn calf serum (Sigma-Aldrich, St. Louis, MO) both performed well and was cost efficient. For ABC169, BM169 was supplemented with 200 µM ascorbic acid (Sigma-Aldrich, St. Louis, MO), 0.2% V/V bovine washed red blood cells (10% suspension, Lampire Biological Laboratories, Pipersville, PA) and 0.2% V/V Porcine cholesterol concentrate (Rocky Mountain Biologicals, Missoula, MT). Ascorbic acid was added from a 500 mM stock dissolved in BM169 that was stored at 4°C in the dark for less than 2 weeks. For regular maintenance, 4-6 worm pairs were cultured at 37°C in 5% CO_2_ in 3 ml media in a 12-well plate; media was changed every other day. To count egg-laying rates, individual worm pairs were maintained in 24 well plates in 1 ml of media and eggs were removed for counting every 24 h during a media change. Worms that separated or became ill during cultivation were excluded from analysis.

### Molecular biology

For quantification of gene expression, RNA was reverse transcribed (iScript, Biorad) and Quantitative PCR was performed using an Applied Biosystems Quantistudio3 instrument and PowerUp SYBR Green Master mix (ThermoFisher, Carlsbad, CA). Gene expression was normalized to the expression of a proteasome subunit (Smp_056500) and relative gene expression values and statistical analyses were performed using the Quantistudio Design and Analysis software (ThermoFisher, Carlsbad, CA). The heatmap depicting relative gene expression was generated using R. Oligonucleotide sequences were listed in S1 table.

### Parasite labeling and imaging

Whole-mount *in situ* hybridization was performed as previously described [26, 54]. ImageJ was used to quantify egg size, autofluorescence, and nuclei number from images of eggs acquired on a Nikon A1+ laser scanning confocal microscope. Like previously described diazo salts [55, 56], Fast Blue BB strongly labeled the polyphenol rich vitelline droplets of mature vitellocytes and could be visualized as both by bright field and fluorescence microscopy. For Fast Blue BB staining, female worms were separated from males using 0.25% tricaine in BM169 and fixed for 4 hrs in 4% Formaldehyde in PBS+0.3% Triton X-100 (PBSTx). Parasites were washed in PBSTx for 10 mins, stained in freshly-made filtered 1% Fast Blue BB in PBSTx for 5 mins and rinsed in PBSTx 3 times. Worms were incubated with 1 µg/ml DAPI in PBSTx for 2hrs, cleared in 80% glycerol in PBS and mounted on slides. For Fast Blue BB/BODIPY 493/503 staining, parasites were processed similarly expect detergents were omitted from all buffers; parasites were labeled with 1 µg/ml BODIPY 493/503 (ThermoFisher, Carlsbad, Ca) in PBS for 1hr following Fast Blue BB staining. For EdU labeling, eggs within 24-48 hours of being laid were incubated in Medium 199 supplemented with 10 µM 5-Ethynyl-2′-deoxyuridine (EdU) and incubated overnight. The following day the eggs were collected by centrifugation at 10,000g for 1min, fixed in 4% formaldehyde in PBSTx for 4hrs, EdU was detected as previously described [54], and washed in PBSTx. *in situ* hybridizations and colormetic detection of Fast Blue BB were imaged using a Zeiss AxioZoom V16 equipped with a transmitted light base and a Zeiss AxioCam 105 Color camera. All other images were acquired using a Nikon A1+ laser scanning confocal microscope.

### Egg culture, miracidia hatching, and labeling

Eggs were collected on 10 µm cell strainer rinsed into Medium 199 (Corning) containing 10% Fetal Bovine Serum and 1x Antibiotic-Antimycotic (Gibco/Life Technologies, Carlsbad, CA) for 7 days at 37°C in 5% CO_2_. After culture, eggs were collected on 10 µm cell strainer and rinsed with artificial pond water 1-2 times and placed in artificial pond water under light. A small aliquot of eggs was counted to determine the total number of eggs examined. Active miracidia were observed and counted under light microscopy every 30 mins for up to 4hrs. Miracidia fixation and labeling was performed as previously described [57].

### Karyotype analysis

Karyotypes from schistosome embryos were determined using a modified version of a previously published method [58]. Briefly, eggs were incubated in M199 for two days after deposition and then incubated in 5 µM Nocodazole for 1 hour at 37°C. Eggs were pelleted at 20,000 g for 1 min, rinsed in DI water, pelleted, resuspended in 1ml of water, and incubated for 20 min at RT. Following centrifugation, the pelleted eggs were resuspended in 1 ml of fixative (3:1 methanol:acetic acid) for 15 minutes. Eggs were again pelleted and resuspended in 100 µl of fixative and the eggshells were disrupted using a Kontes pellet pestle (DWK Life Sciences, Rockwood, TN). The disrupted eggshells were allowed to settle for 10 min, the supernatant was collected, and centrifuged for 8 min at 240 g. All but ∼20 µl of supernatant was removed and the remaining liquid was pipetted dropwise on to a glass slide (Superfrost Plus, ThermoFisher, Carlsbad, CA) freshly dipped in PBS. After air-drying chromosomes were labeled with Vectashield containing DAPI (Vector Laboratories, Burlingame, CA) and imaged on a Nikon A1+ laser scanning confocal microscope with a 60x/1.4 NA objective.

### RNA Interference

Double stranded RNA was generated as previously described [54, 59]. For dsRNA treatment 12 immature female from single-sex infections were soaking in 1 ml Basch 169 supplemented with 1% FBS containing 60 µg/ml of dsRNA for 24 hrs. Then 3 virgin female worms were co-cultured with 3 adult male worms in 3mL ABC169 media with 10% FBS supplemented with 60 µg/ml of dsRNA on day 1, 2, 6 and 10. At D4, unpaired female worms were discarded. At D14, parasites were pulsed for 4 hours with EdU and processed as previously described [54]. Eggs laid between D12 to D14 were collected to determine egg-laying rates.

## ACKNOWLEDGMENTS

We dedicate this manuscript to the memory of Paul Basch. We thank Melanie Cobb, Michael Reese and members of Collins laboratory for comments on the manuscript. Mice and *B. glabrata* snails were provided by the NIAID Schistosomiasis Resource Center of the Biomedical Research Institute (Rockville, MD) through NIH-NIAID Contract HHSN272201000005I for distribution through BEI Resources. The work was supported by the National Institutes of Health (R01 R01AI121037), the Wellcome Trust (107475/Z/15/Z) and the Welch Foundation (I-1948-20180324).

---

**S1 Fig.**
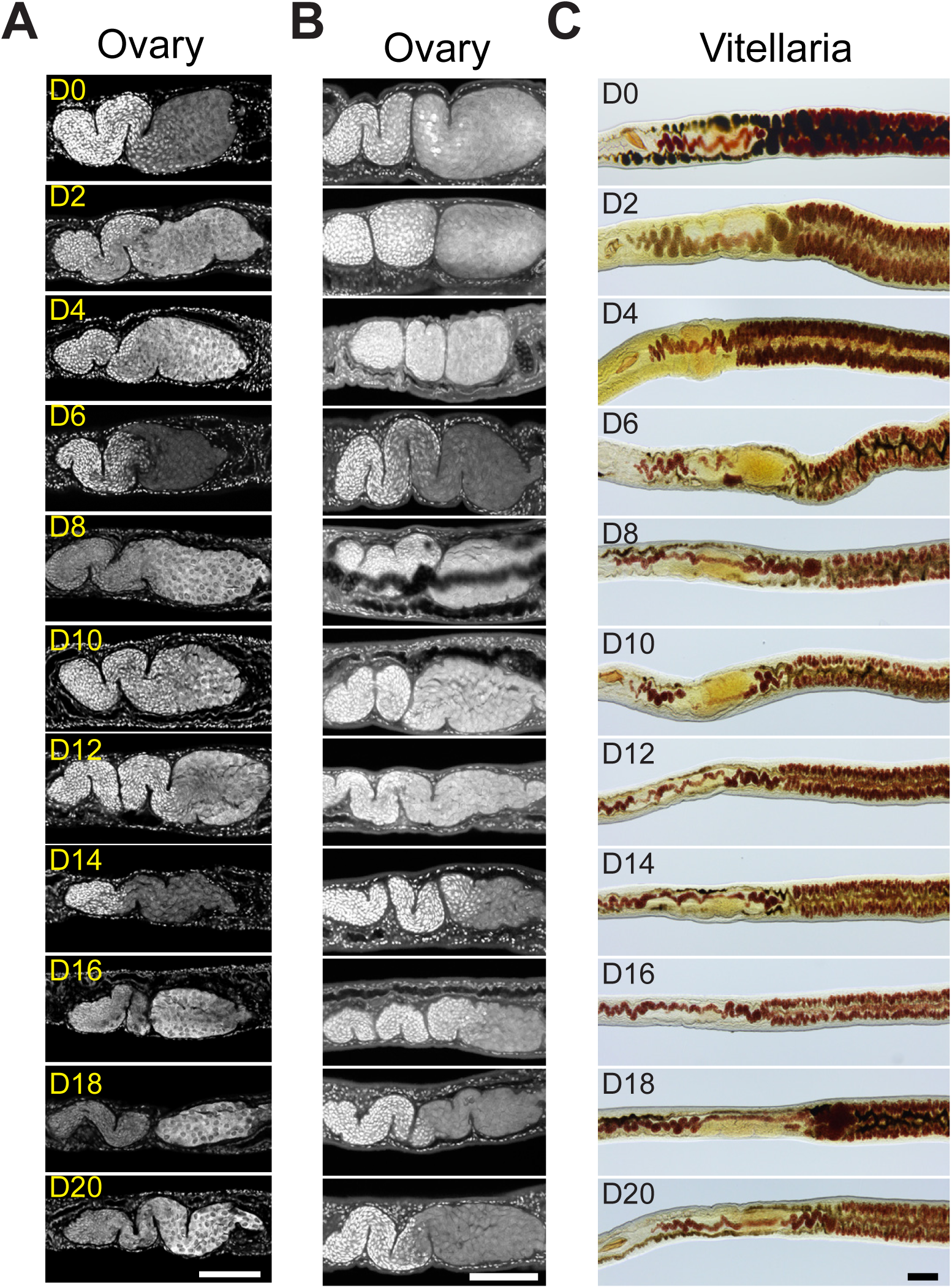
Reproductive changes of paired adult female parasites during *in vitro* culture. Ovaries of females in (A) BM169 or (B) ABC169 between D0 to D20 of in vitro culture labeled with DAPI. Differentiated oocytes are present in the posterior regions (right) of ovaries during in vitro culture regardless of culture condition. (C) Fast Blue BB labeling showing the maintenance of mature vitellocytes during culture in BM169. Representative images from three biological replicates with n > 10 parasites. Scale bars: 100 µm.

**S2 Fig.**
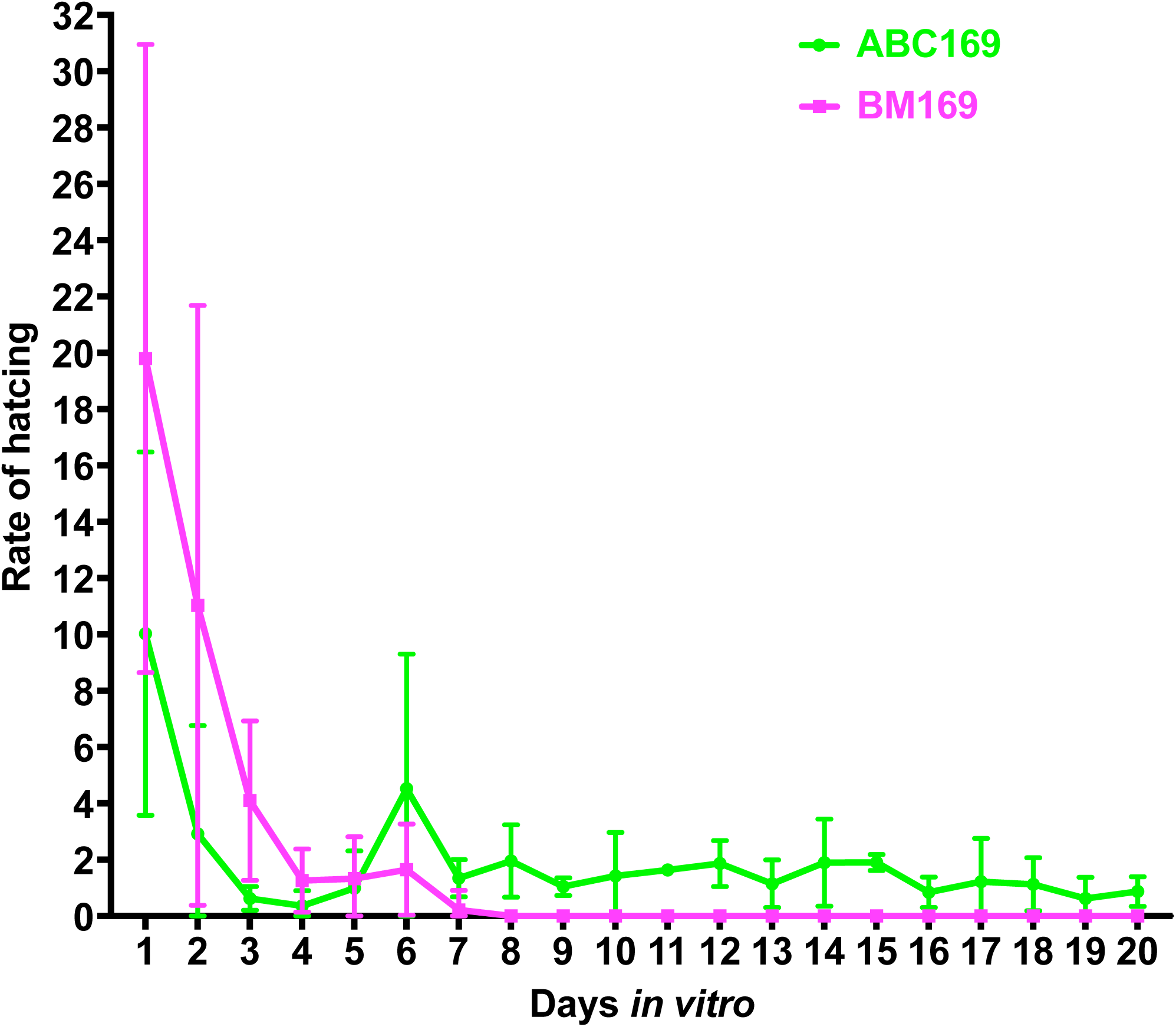
Hatching ratio of eggs laid on different days by worm pairs cultured in BM169 or ABC169. Data points represent mean values from 4 independent experiments. Error bars represent 95% confidence intervals.

**S3 Fig.**
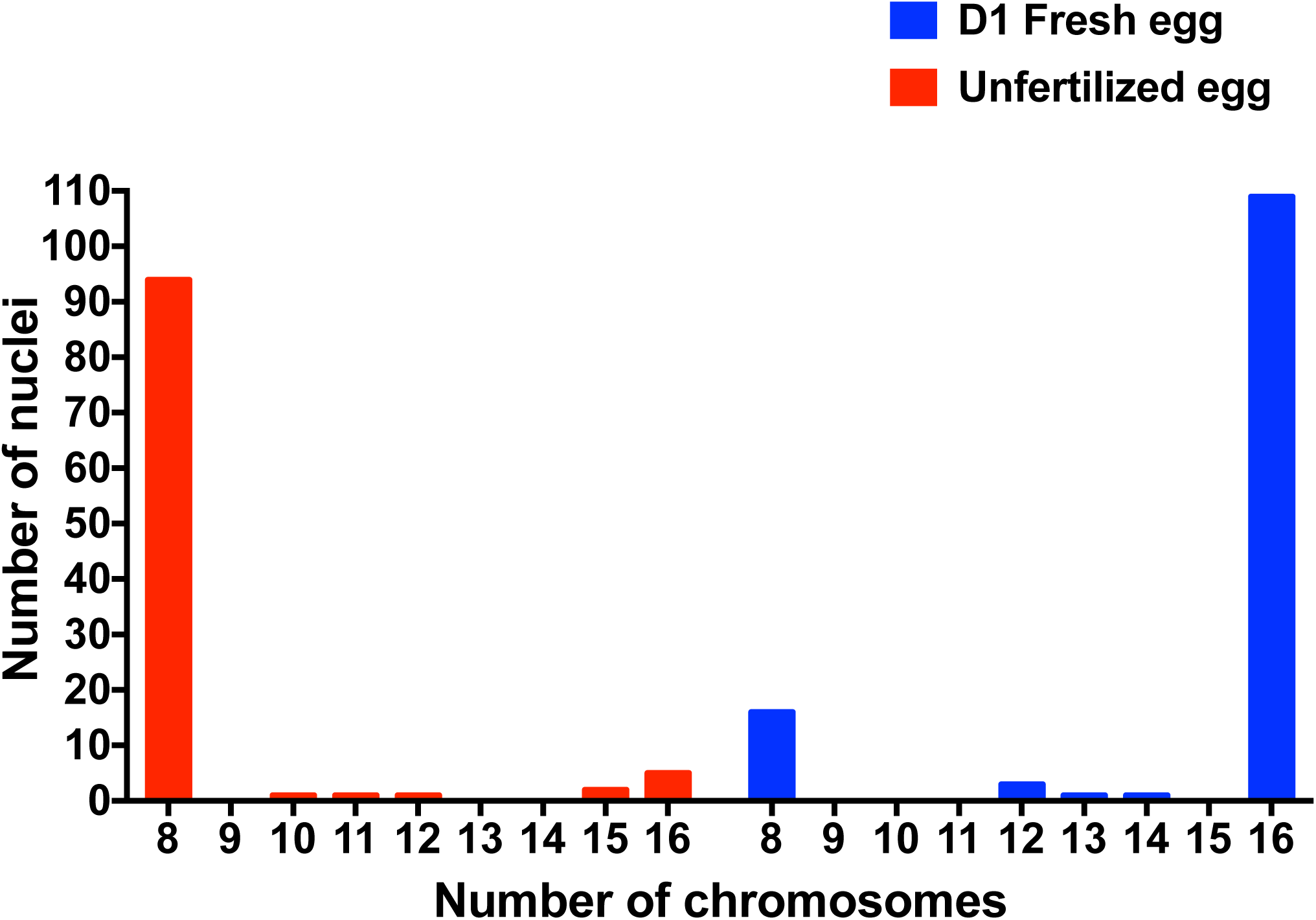
Karyotype analysis on eggs. Plot showing number of chromosomes from karyotypes obtained from eggs laid by freshly perfused worm pairs on D1 of culture (“D1 Fresh eggs”, blue) or females paired with decapitated and castrated male segments (“unfertilized eggs”, red).

**S4 Fig.**
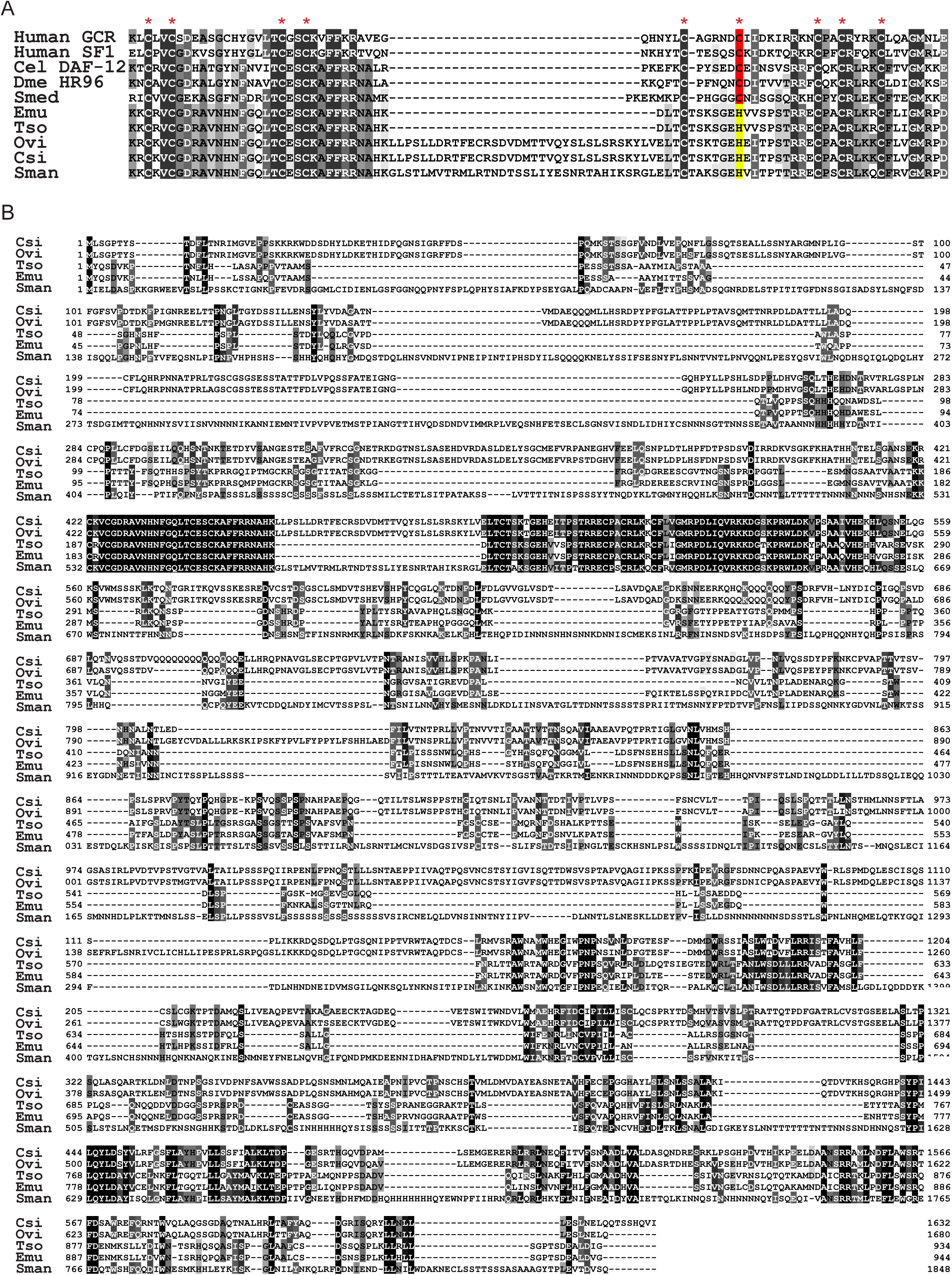
The protein alignment of Smp_248100 and the most similar proteins from other organisms. (A) Sequence alignment showing Smp_248100 shares a conserved DBD with other vertebrate and invertebrate nuclear receptors. Unlike the non-parasitic groups, parasitic flatworms shared a conserved histidine residue (indicted in yellow) in the position of the second conserved cystine (indicted in red) in the second zinc finger. (B) Full-length alignment showing C-terminus of Smp_248100 shares stretches of high amino acid identity with orthologous proteins from other parasitic flatworms. Protein identities: Human GCR, Human glucocorticoid receptor (P04150); Human SF1, Human steroidogenic factor 1(Q13285); Cel DAF-12, *Caenorhabditis elegans* nuclear hormone receptor family member daf-12 (NP_001041239); Dme HR96, *Drosophila melanogaster* hormone receptor-like in 96 (NP_524493); Smed HR96, *Schmidtea mediterranea* (dd_Smed_v6_14067_0_3). Emu, *Echinococcus multilocularis* (EmuJ_001078800.1); Tso, *Taenia solium* (TsM_000102500); Ovi, *Opisthorchis viverrini* (T265_09674); Csi, *Clonorchis sinensis* (csin106676); Sman, *Schistosoma mansoni* (Smp_248100).

**S5 Fig.**
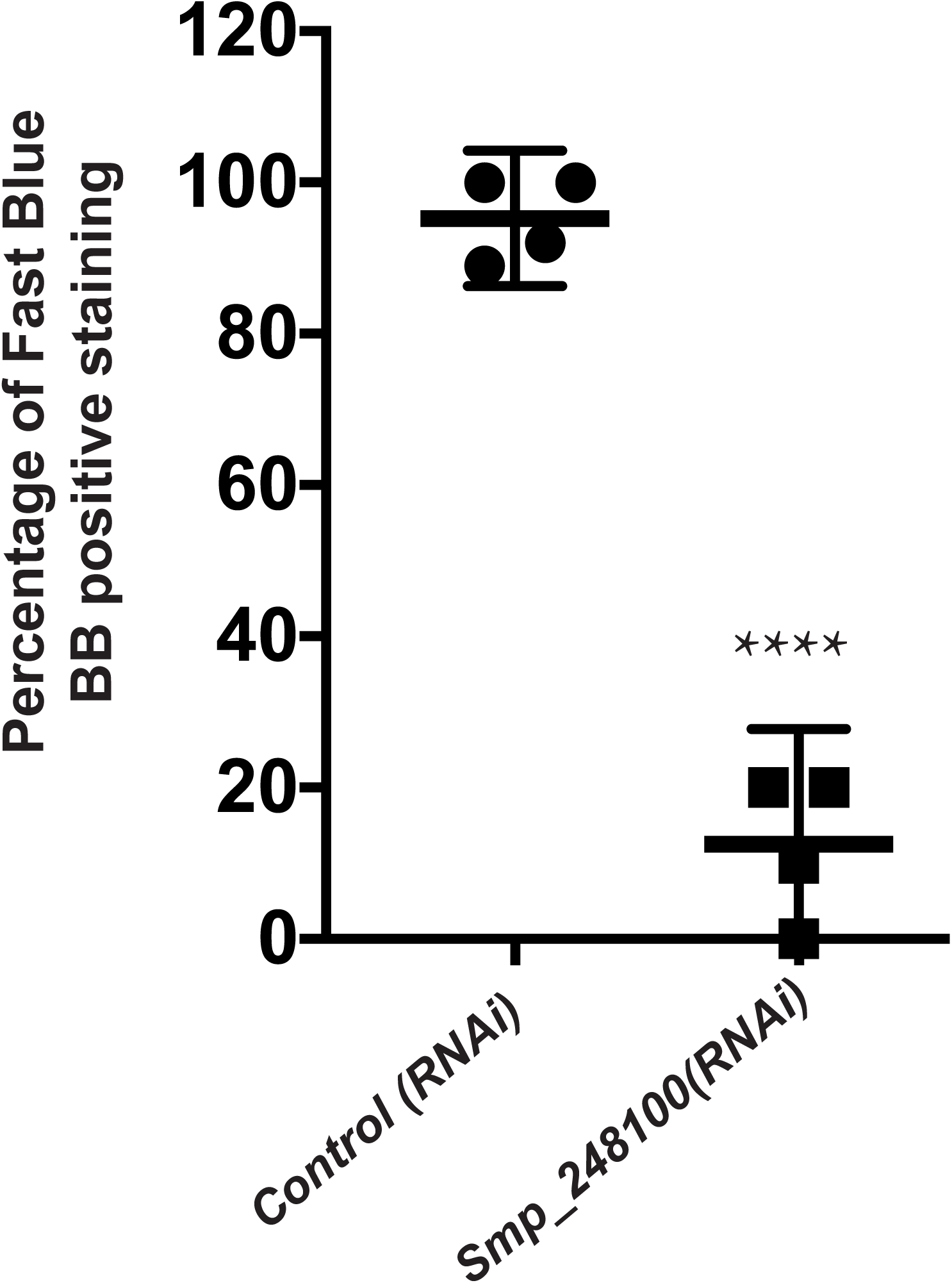
Fast Blue B staining of the dsRNA-treated worms at D14. Plot showing percentage of Fast Blue BB positive labeling in *Smp_248100 (RNAi)* versus *control (RNAi).* Representative images from n > 39 parasites for each treatment examined in 4 separate experiments. **** p<0.0001, T-Test. Error bars represent 95% confidence intervals.

